# Thalamocortical Projectional Divergence Leads to Complexity in Functional Organizations in Mouse Auditory Cortex

**DOI:** 10.1101/226100

**Authors:** Shinpei Ohga, Hiroaki Tsukano, Masao Horie, Hiroki Terashima, Nana Nishio, Yamato Kubota, Kuniyuki Takahashi, Ryuichi Hishida, Hirohide Takebayashi, Katsuei Shibuki

## Abstract

Frequency-related topological projections from the ventral division of the medial geniculate body (MGv) relay the tonotopic organization found in primary auditory cortex (A1). However, relaying circuits of the functional organization to higher-order, secondary auditory field (A2) have not been identified so far. Here, using tracing, we found that A2 receives dense topological projections from MGv in mice, and that tonotopy was established in A2 even when primary fields including A1 were removed. These indicate that thalamic inputs to A2 are sufficient for generating its tonotopy. Moreover, neuronal responses in the thalamocortical recipient layer of A2 showed wider bandwidth and greater heterogeneity of the best frequency distribution than those of A1, which was attributed to larger divergence of thalamocortical projections from MGv to A2 than those from MGv to A1. The current study identifies that the functional organization in the auditory cortex can be determined by the structure of thalamocortical input.

**Significant Statement:** Although peripheral input patterns to the primary auditory cortex (A1) of the brain are well understood, how tonal information is relayed to higher-order regions such as the secondary auditory field (A2) remains unclear. This work revealed a new source of auditory information to A2; the tonal map in mouse A2 is primarily produced by orderly projections from the primary auditory thalamus. We also found that the complex behaviour and organization of neurons in A2 is generated by divergent projections from the primary thalamus that converge on neurons in A2. Our findings indicate that thalamocortical projections constitute a major factor that determines the regional properties and functional organization of mouse A2.

## Introduction

The auditory cortex is composed of multiple regions that form a hierarchy of information processing; tonal information enters at the primary region and is then relayed to higher-order regions in streams of increasing complexity (Kaas and Hackett, 2000). Primary auditory cortex exhibits tonotopy that reflects information directly relayed from topologically connected lemniscal pathways that ascend from the periphery through the ventral division (MGv) of the medial geniculate body (MGB) (Saenz and Langers, 2014). In contrast, many believe that higher-order regions do not show tonotopy because their cortical input is secondarily, and because much of their input from the thalamus originates in the higher-order, non-lemniscal dorsal division (MGd) of MGB, which is not tonotopic. This parallel pathway model, in which one pathway connects MGv to primary regions and the other connects MGd to higher-order regions, has been a basic belief for the last two decades of auditory system research in several animal species, including monkeys (de la Mothe et al., 2012), cats (Huang and Winer, 2000), rats (Smith et al., 2012), and mice (Llano and Sherman, 2008).

The primary auditory cortex (A1) and secondary auditory field (A2) are regarded as representatives of primary and higher-order auditory regions respectively, and exemplify the auditory cortex hierarchy across species (Issa et al., 2014; Lee and Sherman, 2008, 2010; Lee and Winer, 2008). For mice in particular, A1 and A2 are thought to exhibit a cortical hierarchy that is reflected in different response latencies (Guo et al., 2012; Joachimsthaler et al., 2014) and also in different levels of tonotopic complexity in their respective layers 2/3 (Issa et al., 2014). Recently, higher-order regions beyond A1 have been shown to reflect tonotopy in different mammalian species, and thus multiple tonotopic representations are now thought to exist within the auditory cortex (Bizley and King 2009; Harel et al., 2000; Higgins et al., 2010; Kalatsky et al., 2005; Nishimura et al., 2007; Petkov et al., 2006).

Mouse A2 also exhibits tonotopy (Issa et al., 2014; Kubota et al., 2008; Kato et al., 2015), which is presumed to incorporate information relayed from lemniscal regions. However, these theorized topological projections that determine tonotopy in mouse A2 have yet to be identified.

Recent developments in auditory thalamocortical research suggest that tonotopy in A2 might reflect that in the lemniscal thalamus. In canonical theories, A1 and A2 have an intracortical hierarchical relationship in which tonotopy in A2 is derived from secondary connections originating in A1. Thus, mouse A2 should receive frequency-related projections from A1 that are topologically arranged, similar to how mouse primary visual cortex topologically projects to secondary visual cortices (Wang and Burkhalter, 2007; Glickfeld et al., 2013). Another possibility is that tonotopy in A2 is based on input from MGv that reflects its subcortical functional organization. Recent tracing studies suggest that the lemniscal thalamocortical pathway is actually composed of several different parallel pathways. Rather than being a single homogenous structure, MGv is composed of multiple compartments, each of which sends frequency-related topological projections to distinct cortical targets (Horie et al., 2013; Takemoto et al., 2014; Tsukano et al., 2015). At present, the thalamic origins of four cortical tonotopic regions (not including A2) have been identified in rostral and middle parts of MGv (Takemoto et al., 2014; Tsukano et al., 2015). However, the cortical target of neurons located in the caudal portion of MGv remains unknown. Although a detailed morphological study has reported that MGd terminates in A2 of mice (Llano and Sherman, 2008), the authors did not investigate whether thalamocortical inputs to A2 also originate from MGv. Therefore, A2 might receive topologically arranged auditory input from the caudal compartment of the MGv.

Differences in regional properties between A1 and A2 can be seen in tonotopic organization at the single neuron level, in which neighboring neurons are heterogeneously tuned to dissimilar best frequencies (BFs) (Bandyopadhyay et al., 2010; Castro and Kandler, 2010; Rothschild et al., 2010; Tao et al., 2017). Specifically, the extent of heterogeneous tuning within layer 2/3 follows a hierarchy, becoming larger in A2 than in A1 (Issa et al., 2014). Although the origin of this region-specific functional organization is unclear, it is likely formed by a mechanism that depends on either exogenous input or intrinsic local circuits. However, observation of layer 2/3 neurons is hardly sufficient to evaluate these two possibilities. We hypothesized that the heterogeneity of neighboring BFs in thalamo-recipient layers is derived from exogenous input patterns that originate in the caudal part of MGv. If this is true, then it should reflect the degree of anatomical divergence in thalamocortical projections (Imaizumi and Lee, 2014; Storace et al., 2011), which leads to locally heterogeneous frequency inputs when high (Vasquez-Lopez et al., 2017). Investigating any parallel thalamocortical streams that target A1 and A2 will provide an experimental platform for evaluating this hypothesis.

In the current study, we show that the major thalamic origin of mouse A2 is the caudal part of MGv (Figure 8). A2 and caudal MGv are topologically connected, and tonotopy in A2 can be established via thalamic inputs in the absence of corticocortical inputs from A1. Thalamocortical projections from MGv to A2 diverge more than they do to A1, and the tonotopy of the thalamocortical recipient layer 3b/4 is thus more heterogeneous in A2 than in A1.

## Results

### Mouse A2 receives major inputs from the primary thalamic division, MGv

We identified the precise location of right auditory cortical regions using flavoprotein fluorescence imaging (Figure 1A). Tonal stimuli around 5 kHz are known to produce neuronal responses in three regions. Numerous studies of mouse auditory cortex consider the dorsal two regions as the anterior auditory field (AAF) and A1, and the ventral region as A2 (Figure 1B) (Issa et al., 2014; Guo et al., 2012; Joachimsthaler et al., 2014; Kato et al., 2015; Tsukano et al., 2015; Stiebler et al., 1997; Tsukano et al., 2016). In addition to these three areas, the dorsomedial field (DM) was also activated (Figure 1B). Quantitative analysis of A2 using 5 kHz, 30 kHz, and 60 kHz tones indicated that the peak fluorescence change shifted from dorsal to ventral with increased tonal frequency (Figures 1C and 1D), as previously reported (Issa et al., 2014; Kubota et al., 2008; Kato et al., 2015). To investigate the origin of thalamic input to A2, we injected retrograde tracers into A2 after regional identification using optical imaging. We triple injected Fast Blue, CTB-488, and CTB-555 (Takemoto et al., 2014) along the tonotopic axis of A2 (Figures 1E and 1F), prepared consecutive coronal sections, and then visualized fluorescence-positive thalamic neurons.

**Figure 1.**
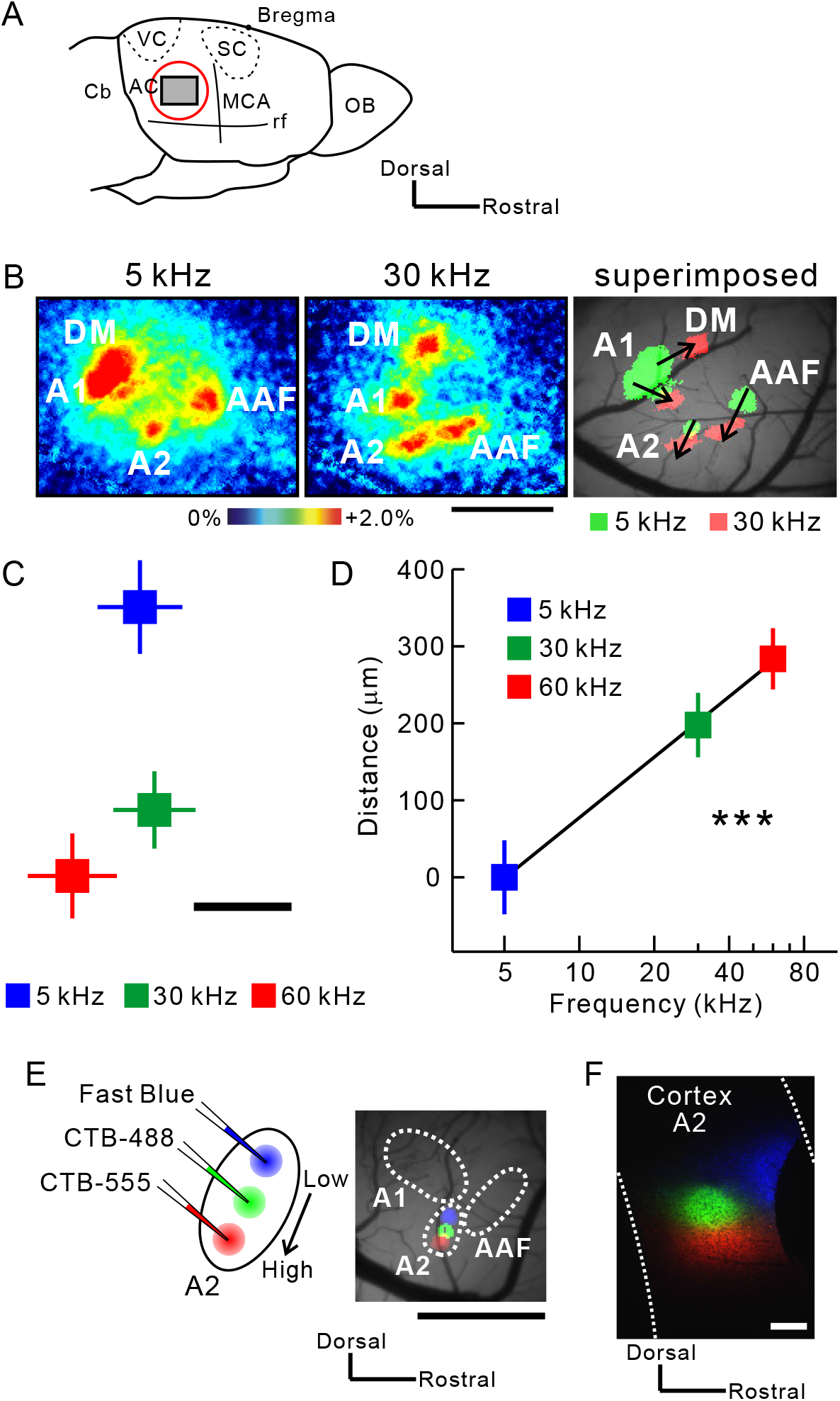
Injection of retrograde tracers along the tonotopic map in mouse A2. (A) A schematic drawing of the moue auditory cortex. AC, auditory cortex; Cb, cerebellum; MCA, medial cerebral artery; OB, olfactory bulb; rf, rhinal fissure; SC, somatosensory cortex; VC, visual cortex. (B) Typical images of tonal responses in the auditory cortex revealed using flavoprotein fluorescence imaging. Scale bar, 1 mm. (C) The tonotopic organization of A2. Peak fluorescence change in response to different frequencies is plotted. Significant shifts in response were observed from dorsal to ventral A2. Data were obtained from 9 mice and superimposed based on the pixel of an A1 5-kHz response peak. Scale bar, 100 μm. (D) Correlation between frequencies and distances. Tonotopy was arranged along the axis connecting the peak pixels for 5 kHz and 60 kHz tones. r = 0.66, \*\*\**p* < 0.001, Spearman’s test; n = 9 each. (E) Triple injection of Fast Blue (blue), CTB-488 (green), and CTB-555 (red) along the tonotopic gradient of A2. Tracer deposits on the brain surface ~30 min after injection are shown on the right. Scale bar, 1 mm. (F) Tracer deposits in A2 on a coronal slice. Scale bar, 200 μm. Data are shown in mean ± SEM.

The boundary between MGv and MGd can be determined based on immunoreactivity (Lu et al., 2009). Here, we used SMI-32 immunolabeling that strongly stains MGv but not MGd (Figure 2A) (Horie et al., 2013; Tsukano et al., 2015), and calbindin immunolabeling that strongly stains MGd but not MGv (Takemoto et al., 2014; Lu et al., 2009). Although some fluorescence-positive neurons were located in MGd as reported previously (Llano and Sherman, 2008), more were found within MGv (Figure 2B). After injecting retrograde neuronal tracers into A2, 43 coronal slices were analyzed. Among the labeled neurons, 63.7% (4,614/7,238 neurons) were found in MGv, 18.7% (1,351/7,238 neurons) in MGd, and 17.6% (1,273/7,238 neurons) in the rest of the auditory thalamus. We next quantitatively analyzed the rostro-caudal distribution of projection neurons in MGv. Neurons projecting to A2 were primarily located between 3.3 and 3.6 mm posterior to the bregma (Figure 3A). AAF and A1 are known to receive thalamic input from the middle part of MGv, around 2.9 to 3.2 mm posterior to the bregma (Horie et al., 2013; Takemoto et al., 2014; Tsukano et al., 2017). We found that neurons projecting to A2 were clearly located caudal to those projecting to AAF/A1 (Figure 3B). We directly confirmed the rostrocaudal locations of thalamic neurons by preparing horizontal slices (Figure 3C). These results clearly indicated that mouse A2 receives lemniscal projections originating from the caudal part of the MGv.

**Figure 2.**
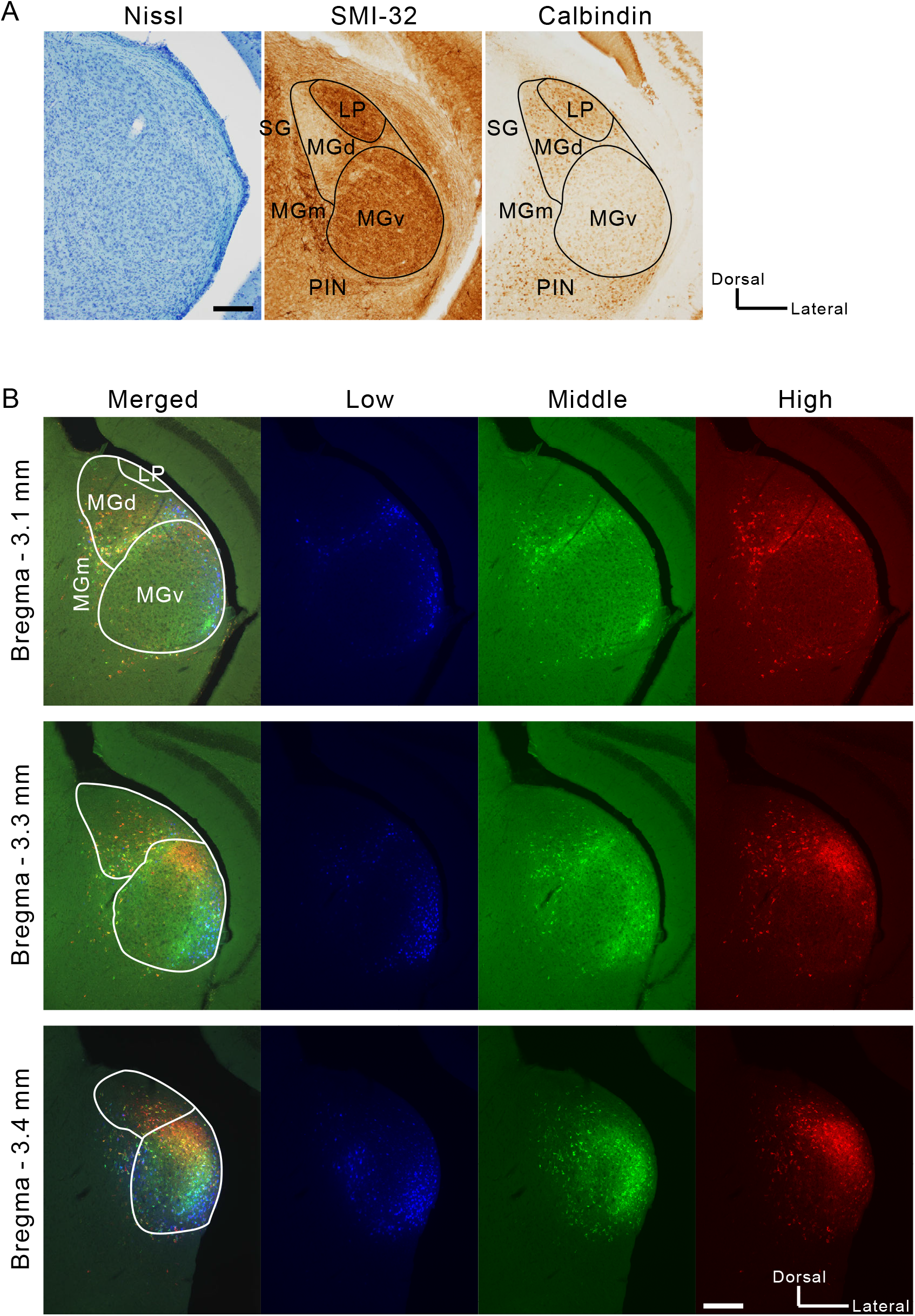
Visualizing thalamic neurons that project to A2. (A) An example pair of Nissl staining and immunolabeling in slices ~3.0 mm posterior to the bregma. These images were obtained from a single animal. LP, lateral posterior nucleus; PIN, posterior intralaminar nucleus; SG, suprageniculate nucleus. Scale bar, 200 μm. (B) Distribution of neurons that were labeled with tracers injected into A2. Blue, green, and red neurons give rise to projections to low, middle, high frequency areas in A2. Scale bar, 200 μm.

**Figure 3.**
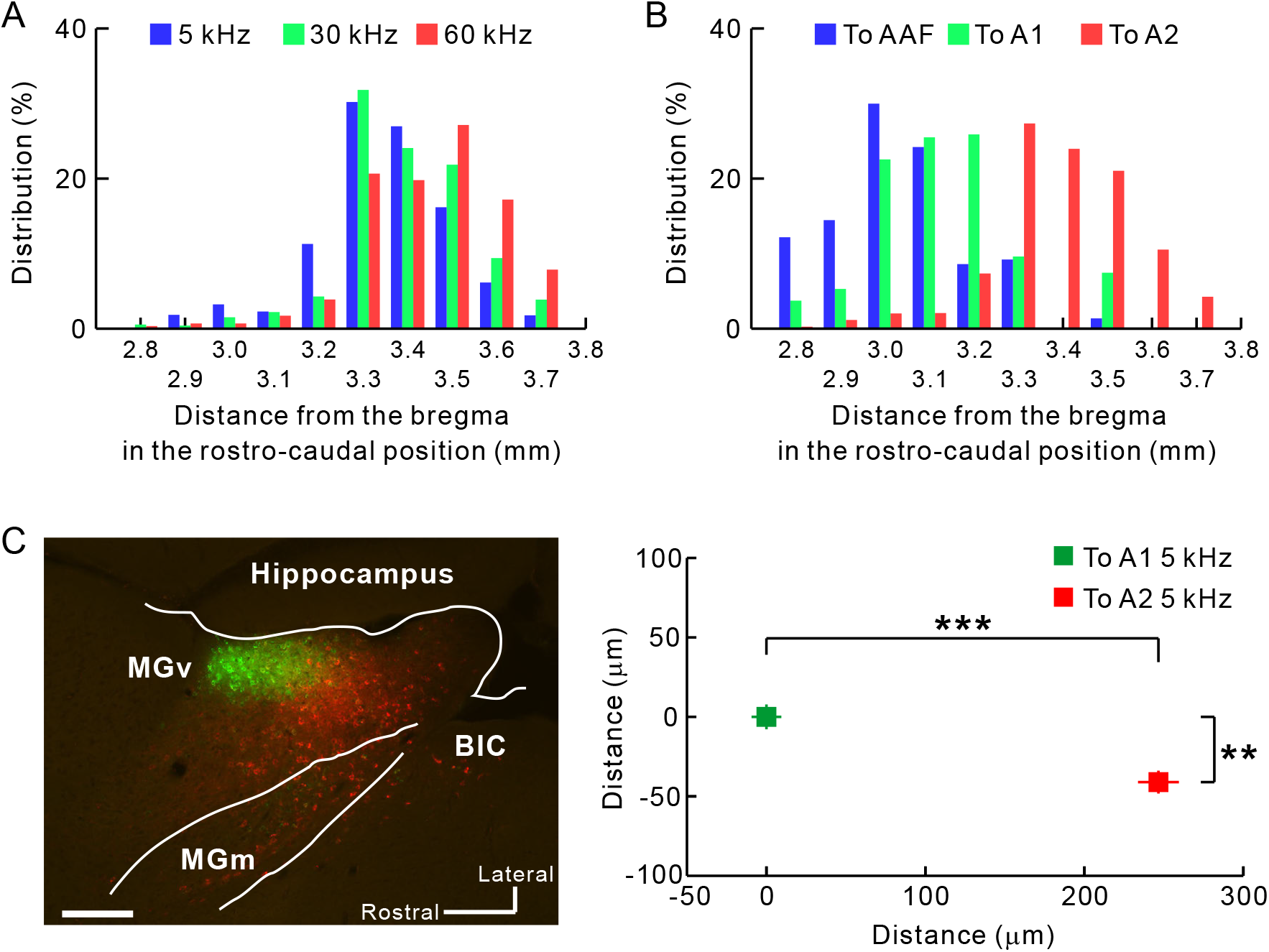
Quantification of the distribution of MGv neurons projecting to A2. Rostrocaudal distribution of thalamic neurons projecting from MGv to A2. Averaged coordinates were 3.34 mm, 3.39 mm, and 3.43 mm posterior to the bregma for low, middle, and high frequencies, respectively. 5 kHz vs. 30 kHz, *p* < 10^−6^; 5 kHz vs. 60 kHz, *p* < 10^−35^; 30 kHz vs. 60 kHz, *p* < 10^−5^, *χ*^2^ test after Bonferroni correction. (B) The caudal part of MGv as a thalamic origin of A2. Data in blue, green, and red indicate distribution of neurons projecting to AAF, A1, and A2, respectively. Averaged coordinates were 3.03 mm, 3.13 mm, and 3.93 mm posterior to the bregma for AAF, A1, and A2, respectively. AAF vs. A1, *p* < 10^−35^; AAF vs. A2, *p* < 10^−35^; A1 vs. A2, *p* < 10^−35^, *χ*^2^ test after Bonferroni correction. To AAF, n = 1,314 cells obtained using fluorescent CTB (5 mice); to A1, n = 510 cells obtained using fluorescent CTB (5 mice); To A2, n = 2,718 cells used in (A) and (B). (C) Confirmation of caudal outputs to A2 using horizontal slices. Neurons projecting to A2 (red) were located significantly caudally compared with those projecting to A1 (green) (\*\**p* < 0.0001; \*\*\**p* < 10^−34^, Mann–Whitney U-test; n = 196 neurons projecting to A1; n = 198 neurons projecting to A2; from 3 mice). BiC, brachial nucleus of the inferior colliculus. Scale bar, 200 μm. Data are shown in mean ± SEM.

Next, we investigated whether neurons projecting to A2 are arranged in a frequency-related fashion. Using coronal slices, we obtained the relative location of each fluorescence-positive neuron in reference to the MGv center (Figure 4A). MGv neurons projecting to the low, middle, and high frequency areas of A2 were topologically arranged in the ventrodorsal direction of caudal MGv (Figures 4B and 4C). Comparing plots between Figures 4B and 4C, we found that overall positions shifted ventrally in caudal slices; therefore, we evaluated frequency-related positional shifts using horizontal slices. We injected CTB-555 and CTB-488 into high and low frequency areas of A2. In the dorsal part of MGv, almost all neurons projected to high frequency areas (Figure 4D). Moving ventrally, neurons projecting to low frequency areas began to appear rostral to those projecting to high frequency areas (Figure 4E). Going more ventrally, neurons projecting to high and low frequency areas were evenly distributed (Figure 4F). In the ventral part, we found that neurons projecting to high frequency areas were shoved caudally (Figure 4G). These data suggest that the caudal compartment of MGv has a distinct frequency-related structure, and that the gradient is arranged largely along a ventrorostral to dorsocaudal axis, as shown in the parasagittal illustration in Figure 4H. This architecture is consistent with the findings obtained from the coronal slices, which showed that MGv neurons projecting to higher frequency areas of A2 tend to be located more caudally (Figure 3A).

**Figure 4.**
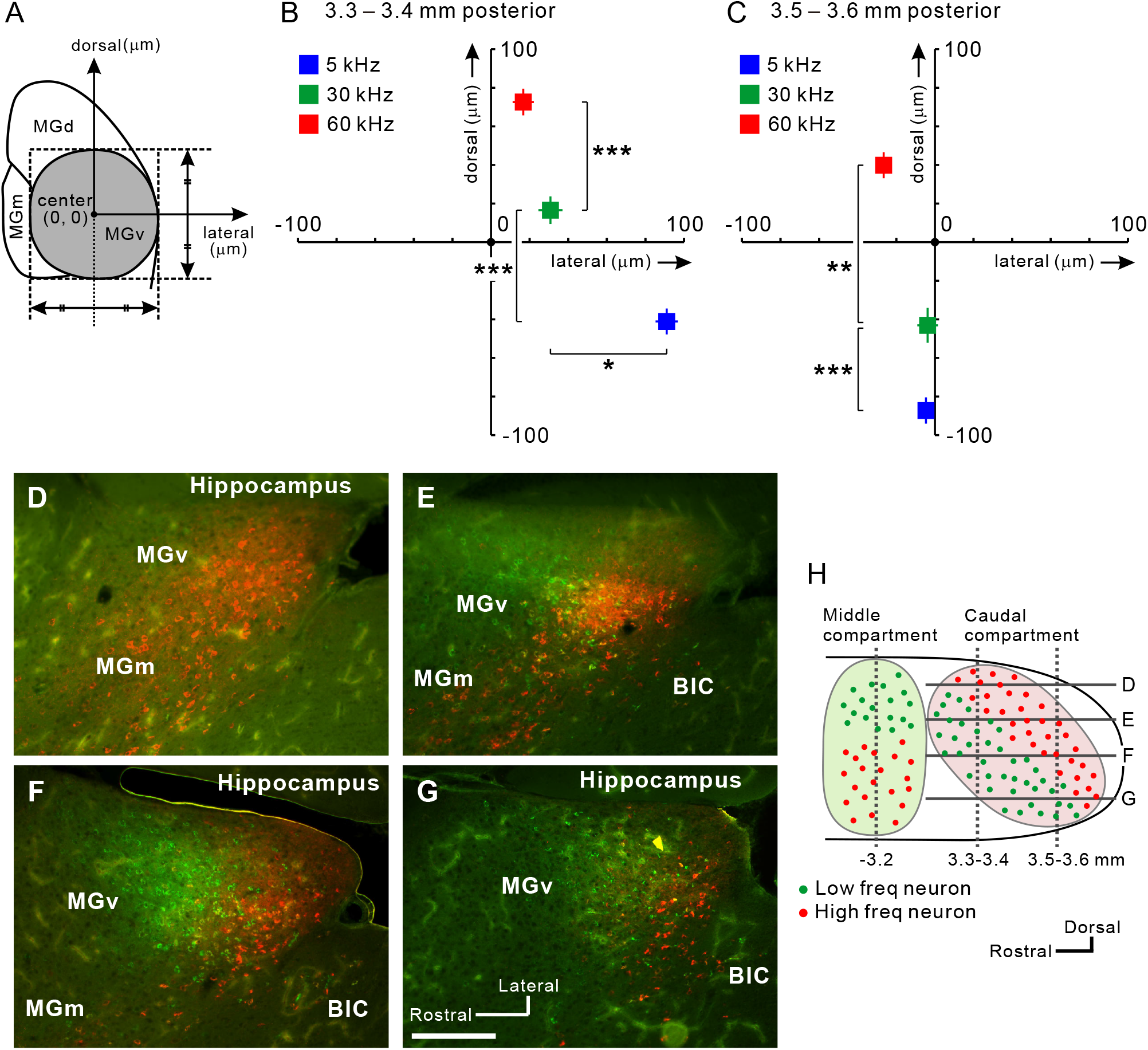
Frequency-related gradient traveling ventrodorsally in caudal MGv. (A) Definition of the MGv center in each slice. (B) Average coordinates for projection neurons to A2 in slices 3.3–3.4 mm posterior to the bregma. The number of neurons was 901 for 5 kHz, 404 for 30 kHz, and 468 for 60 kHz. \**p* < 0.05, \*\*\**p* < 10^−10^, Mann–Whitney U-test. (C) Average coordinates for projection neurons to A2 in slices 3.5–3.6 mm posterior to the bregma. The number of neurons was 286 for 5 kHz, 226 for 30 kHz, and 511 for 60 kHz. \*\**p* < 0.0001, \*\*\**p* < 10^−10^, Mann–Whitney U-test. (D–H) View of the caudal compartment of MGv. Horizontal distribution of MGv neurons projecting to low (green) and high (red) frequency areas in A2 are shown in (D-G). Scale bar, 200 μ m. Approximate dorsoventral levels of each image are indicated in (H). Data are shown in mean ± SEM.

### Frequency-related organization in higher-order auditory thalamus, MGd

Although MGv turned out to be the major origin of thalamic projections to A2, almost 20% of projections originated in MGd. Triple tracer injection revealed that MGd neurons between 3.0 and 3.4 mm posterior to the bregma projected to A2, and that, for tonal stimuli between 5 and 60 kHz, the rostrocaudal distributions of neurons that project to each part of A2 did not differ (Figure S1A). After injecting retrograde tracers into A2, the number of labeled neurons in MGd was small and few slices exhibited neurons that were labeled by all three tracers; however, we did observe positional shifts of labeled neurons (Figure S1B). We quantitatively analyzed the locations of labeled neurons by determining the center of MGd as the origin (Figure S1C) and found that positions shifted from dorsomedial to ventrolateral MGd as the BFs of the target region in A2 shifted from high to low. To the best of our knowledge, this is the first study to demonstrate the existence of frequency-based organization in rodent MGd.

### Tonotopy of A2 is established via thalamic input

A2 is a higher-order region that receives auditory information from A1. However, the results given above showed that A2 also receives distinct topological thalamic inputs. This suggested that tonotopic organization in A2 might exist even when it is isolated from primary cortical regions. To test this hypothesis, we prepared mice in which all auditory cortical regions except for A2 were removed by suction (Figure 5A). A2 in these mice still exhibited robust tonal responses (Figure 5B). While no tonal responses were detected from the removed regions, in A2 they remained even after suction, albeit they were half as intense (Figures 5C and 5D), suggesting that about half of each response was derived corticocortically and half thalamocortically. Importantly, the tonotopic structure of A2 remained after destroying the surrounding cortical regions (Figures 5E and 5F). Moreover, we verified that functional organization in the auditory cortex and response amplitude elicited by amplitude-modulated pure tones was not affected by total removal of the contralateral auditory cortex (Figure S2). This might have resulted from callosal connections that help unify auditory space (Gazzaniga, 2000) and are not necessary for simple tonal processing. These data indicate that parallel topological thalamocortical pathways from MGv give rise to the different tonotopies seen in A2 and A1 (Figure 8).

**Figure 5.**
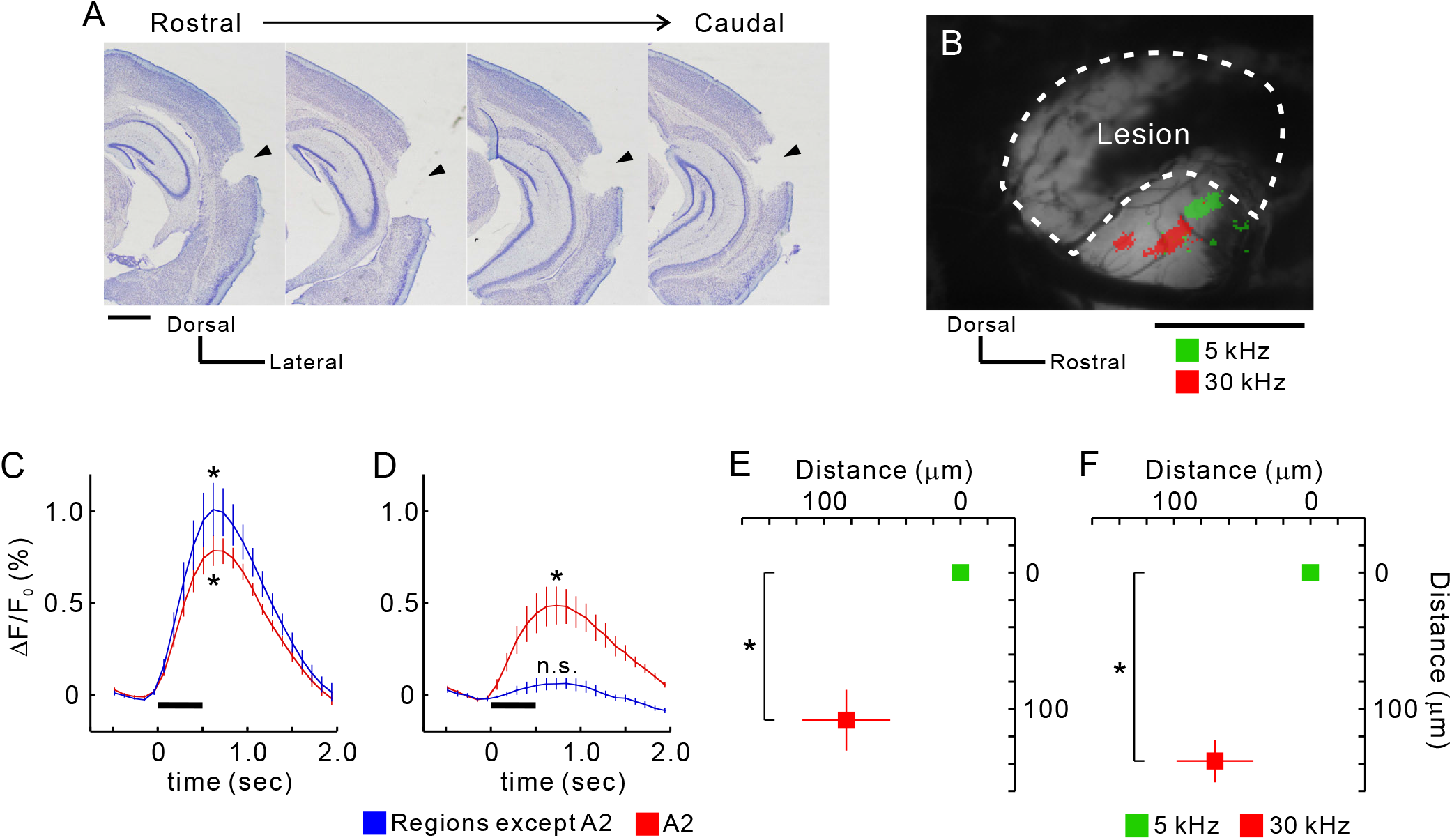
Retained tonotopy in A2 after removal of other auditory regions. (A) Coronal slices from the right hemisphere after suction of auditory cortex that excluded A2. Lesion sites are indicated by arrow heads. Scale bar, 1 mm. (B) Example tonal responses in A2 after suction. Areas enclosed by dotted lines were removed and the white matter on the hippocampus was exposed. Scale bar, 1 mm. (C and D) Temporal profiles of tonal responses before (C) and after suction (D). Data from regions other than A2 are average values from AAF, A1, and DM. Peak values were significantly larger than baseline values. \**p* < 0.05, Wilcoxon signed-rank test. Four profiles were obtained from the same animals (n = 5 mice). (E and F) Quantitative analysis of tonotopic shifts before (E) and after suction (F). Coordinates were averaged based on the peak pixels of 5 kHz areas. \**p* < 0.05, Wilcoxon signed-rank test. Four data were obtained from the same animals (n = 5 mice). Data are shown in mean ± SEM.

### Heterogeneous tonotopy of A2 reflects divergent thalamocortical projections

Previous tracing studies have shown clear topological projections from MGv to A1 that have little spatial overlap between the two middle MGv neuronal populations that project to high and low frequency areas of A1 (Horie et al., 2013; Takemoto et al., 2014). In contrast, caudal MGv neurons that project to A2 exhibited seemingly overlapping distributions, although they also showed a ventrodorsal tonotopic gradient at the macroscopic level (Figure 2). To quantify and compare the degree of spatial overlap in the MGv neuronal populations, we dual-injected CTB-555 and CTB-488 into high and low frequency areas of A1 and A2, respectively (Figure 6A). An analysis of spatial spread in four directions showed that neurons projecting to A2 were distributed more widely than those projecting to A1 in the dorsoventral direction, which corresponds to the tonotopic axis (Figure 6B). As a result, the separation index along the dorsoventral axis was significantly smaller for neurons projecting to A2 than those projecting to A1 (Figure 6C). These data suggest that neurons in the middle part of MGv and neurons in the thalamocortical recipient layer of A1 are connected with a strictly topological relationship. In contrast, neurons in the caudal part of MGv project to a broader area of A2 with a looser topological relationship (Figure 6D).

**Figure 6.**
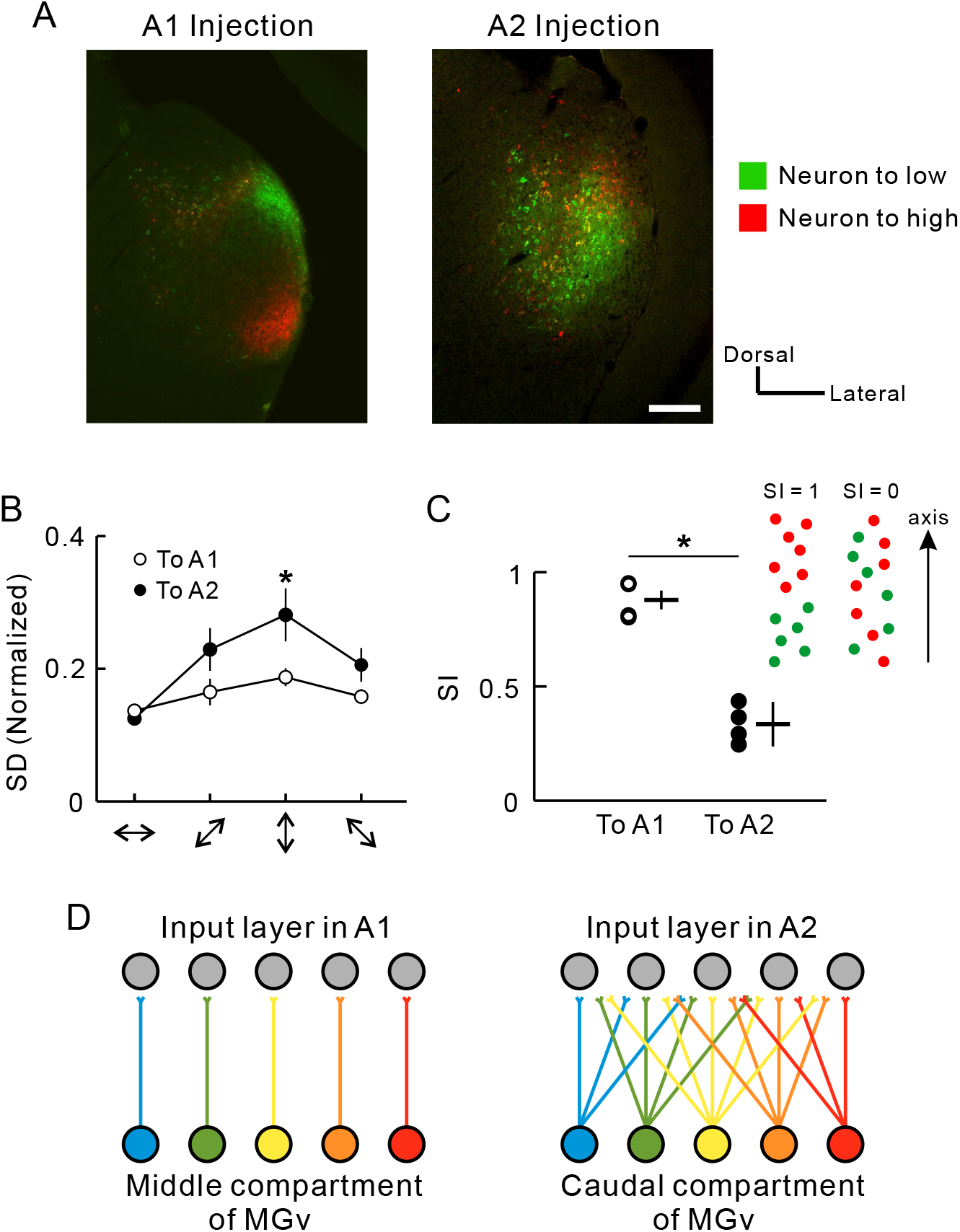
Divergence of thalamocortical projections from MGv to A2. (A) Examples of MGv neurons projecting A1 or A2. Scale bar, 200 μm. Green, CTB-488 that was injected into a cortical low frequency area of 5 kHz. Red, CTB-555 that was injected into a cortical high frequency area of 60 kHz. The right image was modified in brightness. Scale bar, 200 μm. (B) Standard deviations (SD) of the distribution of CTB-positive neurons in four orientations. SD was compensated by width and length of MGv. n = 8 slices from 4 mice. \**p* < 0.05, Mann–Whitney U-test. (C) The separation index (SI) that evaluates extent of how intermingled CTB 488-positive and CTB 555-positive neurons are. SI ranges from 0 to 1, with one meaning total separation (see Methods). Data were obtained from four slices for each region. \**p* < 0.05, Mann–Whitney U-test. (D) Summary illustration of Figure 6. Data are shown in mean ± SEM.

Studies have suggested that BF spatial distribution becomes more heterogeneous the further a region is along in the hierarchy away from primary cortex (Issa et al., 2014; Winkowski and Kanold, 2013). We expected that neurons in the thalamocortical recipient layer of A2 might receive a wide range of information from broad areas in MGv because of the divergence in thalamocortical projections from MGv. We evaluated this effect by looking at bandwidth and heterogeneity of the BF spatial distribution in the thalamocortical recipient layer using two-photon calcium imaging. Thalamocortical recipient layers in the auditory cortex are a deeper part of layer 3 and layer 4 (Huang and Winer, 2000). They are equivalent to layers 3b/4 in mice, which are located ~300–450 mm beneath the cortical surface (Winkowski and Kanold, 2013; Oviedo et al., 2010; Zhou et al., 2014). We observed tonal responses of neuronal cell bodies 377 ± 41 μm (mean ± SD) beneath the cortical surface (Figure 7A). Neurons in A1 tended to respond to a specific tonal stimulus. In contrast, neurons in A2 had a tendency to respond to a wider frequency range (Figure 7B). Analysis showed that neuronal bandwidth was significantly larger in A2 than in A1 (Figure 7C), which suggests that A2 neurons are more broadly tuned than A1 neurons because of convergent input from caudal MGv (Figure 6D). Next, we investigated differences in BF spatial distribution between A1 and A2. Although a study has suggested that local BF spatial distribution in the recipient layer of A1 was totally homogeneous (Winkowski and Kanold, 2013), here we found that it was somewhat heterogeneously arranged (Figure 7D). Even so, the BF and distance along the tonotopic axis were significantly correlated in A1 (r = 0.61; Figure 7E), as were the difference in BFs and the distance between pairs of neurons (r = 0.43; Figure 7F). In contrast, the spatial distribution of BFs was coarse in A2. Although the BF and distance along the tonotopic axis were significantly correlated (r = 0.30; Figures 7G and 7H), the correlation was much weaker than what we observed for A1. Similarly, although we found a significant correlation between the difference in BFs and the distance between pairs of neurons, the correlation coefficient (r = 0.05) was clearly smaller than that for A1 (Figure 7I). Taken together, these data suggest that the anatomical divergence of thalamocortical projections from MGv is reflected in the functional organization of the thalamocortical recipient layer in the auditory cortex.

**Figure 7.**
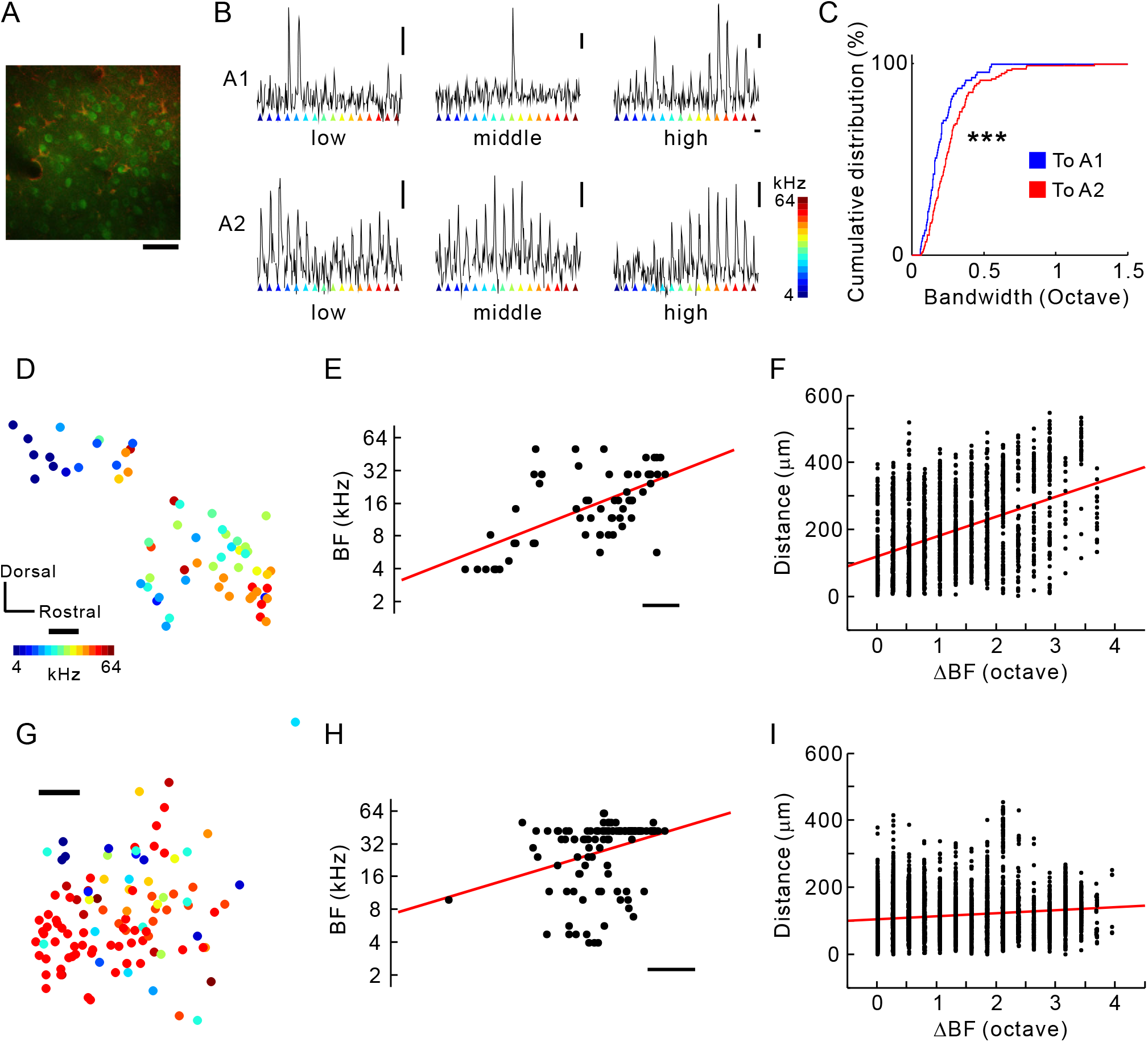
Complex responses of neurons in the thalamocortical recipient layer of A2. (A) An example plane of two-photon imaging. Cal-520-positive neurons and SR-101-positive astrocytes are shown in green and red, respectively. Scale bar, 50 μm. (B) Responses of a low, middle, or high frequency-responsive neuron in A1 (top), and those of a low, middle, or high frequency-responsive neuron in A2 (bottom). Vertical bar, 10%; horizontal bar, 3 s. (C) Cumulative distribution of neuronal bandwidth in A1 and A2. \*\*\**p* < 10^−10^, Kolmogorov-Smirnov test. (D and E) Distribution of best frequencies (BF) in A1. A significant relationship was found between distance and best frequencies (BFs) (n = 62 neurons from two mice; r = 0.61, *p* < 0.0001, Spearman’s test). Scale bar, 50 μm. (F) Relationship between differences in BF and distance (1891 pairs, r = 0.43, *p* < 0.0001, Spearman’s test). (G and H) Distribution of BFs in A2. A significant relationship was observed between distance and BF (n = 109 neurons from two mice; r = 0.30, *p* < 0.01, Spearman’s test). R-values between data in (E) and (H) were significantly different (*p* < 0.05, see Experimental Procedures). Scale bar, 50 μm. (I) Relationship between differences in BF and distance (5,886 pairs, r = 0.15, *p*< 0.0001, Spearman’s test). R-values between data in (F) and (I) were significantly different (*p* < 10^−35^, see Methods).

**Figure 8.**
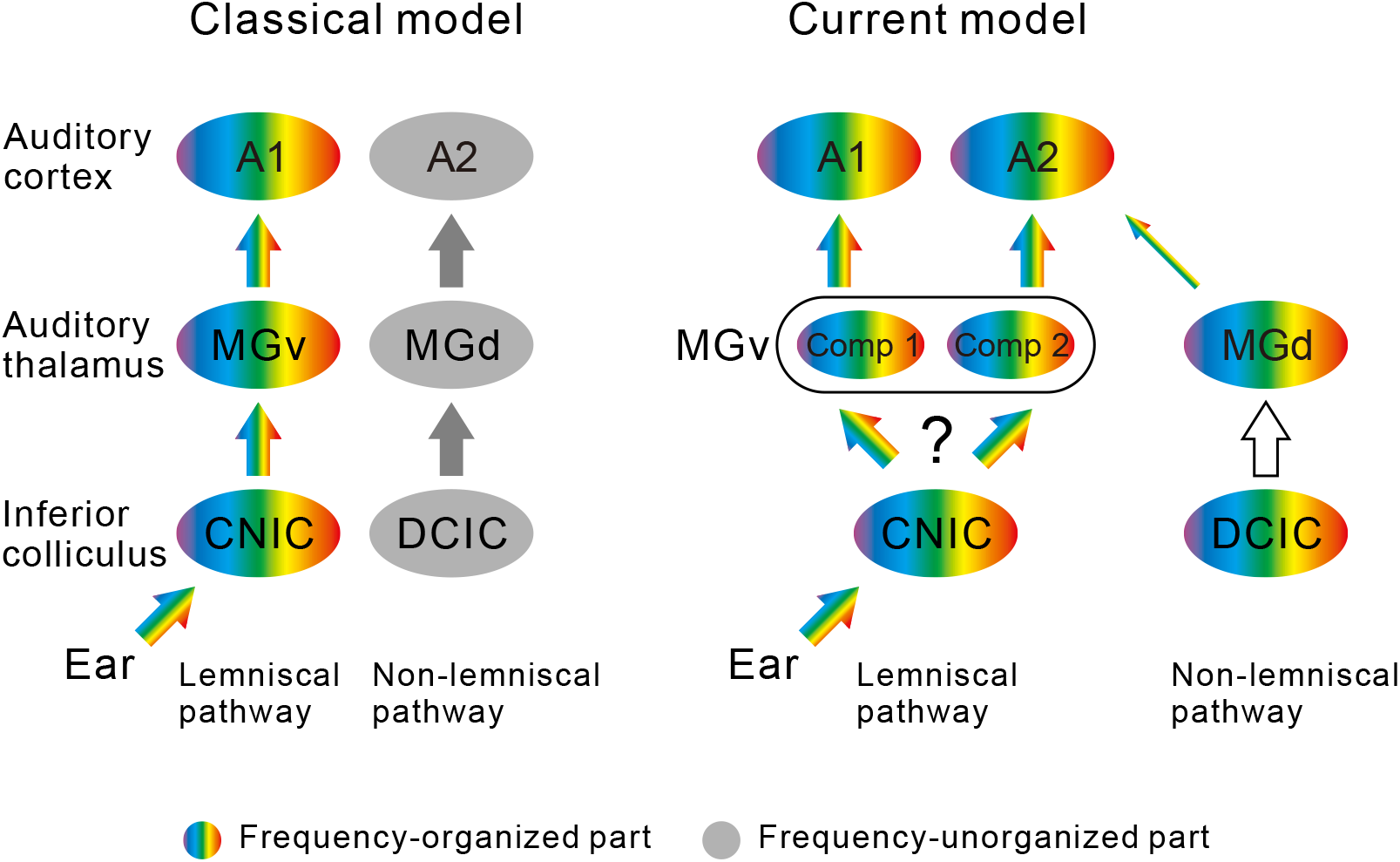
Illustration of a new thalamocortical parallel processing model in the mouse auditory system. The left illustration shows the classical two streams being fed into A1 and A2. These streams correspond to lemniscal and non-lemniscal pathways. The right illustration shows the new scheme revealed in the current study, in which the parallel lemniscal inputs feed into A1 and A2 in mice. Three points have not been clarified; whether divergence of frequency organizations emerges when entering MGB from ICC or not, whether MGd reflects tonotopy, and whether tectothalamic pathways between DCIC and MGd are connected topologically. CNIC, central nucleus of the inferior colliculus (IC); Comp, compartment; DCIC, dorsal cortex of IC.

## Discussion

### Mouse A2 is a higher-order auditory field, but a lemniscal region

In the classical view, the auditory cortex is composed of the core and belt regions (Kaas and Hackett, 2000). By definition, the core comprises the primary regions such as A1/AAF that are centrally localized and have tonotopy that reflects thalamic input from MGv. The belt comprises non-tonotopic, higher-order regions that receive non-lemniscal thalamic inputs from MGd. Mouse A2 has been considered as part of the belt because electrophysiological mapping indicated no clear tonotopic arrangement (Stiebler et al., 1997). However, more recent studies have shown that mouse A2 does indeed have distinct tonotopy (Issa et al., 2014; Kubota et al., 2008), and the current study shows that the tonotopy in A2 also reflects topological thalamic input derived from MGv (Figure 4). This suggests that mouse A2 may belong to the core rather than the belt. To date, we know of four tonotopic regions in the auditory cortex—AAF, A1 A2, and DM (Tsukano et al., 2015)—and one tonotopic region in the insular cortex—the insular auditory field (IAF) (Takemoto et al., 2014). All of these regions receive dense projections from MGv (Tsukano et al., 2017). Such multiple tonotopic fields have been found in rats as well (Storace et al., 2010, 2012), and therefore might be ubiquitous in the rodent auditory system.

The presence of multiple tonotopic regions that receive lemniscal input does not mean that the hierarchy between auditory regions does not exist. Although a study which stimulated sliced auditory cortex suggested bidirectional processing between A1 and A2 (Covic and Sherman, 2011), several in vivo studies suggest that compared with A1, mouse A2 has higher-order properties. For example, latencies for tonal presentation are longer in A2 than in A1 (Guo et al., 2012; Kubota et al., 2008). Additionally, the distribution of BFs for neighboring neurons in layer 2/3 (Issa et al., 2014) and layer 3b/4 (Figure 7) is more mixed in A2 than in A1. Furthermore, as shown here, A2 receives lemniscal thalamocortical projections that are more divergent than those sent to A1 (Figure 6). These findings imply that the nature of A2 is more complex than the primary regions (Joachimsthaler et al., 2014). Moreover, other studies have shown that auditory information propagates unidirectionally even between canonical cores (AAF and A1) (Kubota et al., 2008; Zhang et al., 2017), suggesting that the intracortical serial information processing along a hierarchical stream exists in the auditory cortex. Overall, A2 should still be considered a higher-order region in the mouse auditory cortex, and thus the mouse auditory cortex serves as a good model for investigating serial information processing in the mammalian auditory system.

### Caudal MGv is a thalamic origin of exogenous inputs to A2

The auditory pathway is composed of two major and mutually exclusive canonical parallel streams: the lemniscal and non-lemniscal pathways. The lemniscal pathway conveys auditory information that retains the tonotopic structure that is generated in the ear, and enters AAF/A1 via MGv The non-lemniscal pathway, whose origin is not clear, possibly conveys multi-modal information to A2 via MGd. The current results show that more than half of thalamic projections to A2 originate in MGv, while less than a quarter of them originate in MGd (Figure 3). These data suggest that more information is fed into A2 from MGv than from MGd. Past studies in mice (Llano and Sherman, 2008) and cats (Lee and Winer, 2008) suggested that MGd was the origin of the thalamic inputs to A2. The present results confirm that A2 does receive non-lemniscal thalamic inputs from MGd, but clearly, the region does not have a single source of thalamic input from MGd.

The new scheme of thalamic inputs to A2 found in the present study is compatible with serial information transfer to A2 through the auditory cortical hierarchy. Generally, sensory information transfer to secondary fields is thought to be realized by corticocortical connections. Direct projections from A1 to A2 have been found in mice (Covic and Sherman, 2011) and cats (Lee and Winer, 2008), which are similar to those between primary and secondary fields in the visual (Glickfeld et al., 2013) and somatosensory (Chen et al., 2013) systems. Additionally, auditory information might be conveyed from A1 to A2 indirectly through corticothalamocortical loops, which compose A1-MGd-A2 projections (Lee, 2015). The latter possibility is supported by the presence of projections from A1 to MGd and from MGd to A2 in cats (Huang and Winer, 2000; Winer et al., 1999) and mice (Llano and Sherman, 2008). Our results also confirmed the MGd to A2 pathway in mice (Figure S1). Our current results identified a third and previously unknown pathway that conveys a large amount of tonal information directly from MGv to A2 without passing through A1. Such parallel pathways from the primary thalamus to higher-order auditory fields have been found in rats (Storace et al., 2011; Shiramatsu et al., 2016), where cortical regional properties inherit properties of terminating thalamocortical axons (Storace et al., 2012). Integration of these three types of input in A2 is thought to be critical for realizing higher-order auditory function.

The compartmentalization of rodent MGv has been verified by recent studies as well as the current one. In cats, the lemniscal thalamocortical streams that connect MGv and auditory cortex are parallel because few MGv neurons project to both AAF and A1, even though the two neuronal groups projecting to AAF and A1 are not spatially segregated (Lee et al., 2004; Lee and Sherman, 2010). Recent studies have suggested that mice have much clearer localization and compartmentalization of neuronal groups within MGv that are based on cortical target. Neurons projecting to DM, AAF, or A1 are localized in the rostral, medio-medial, and medio-lateral compartments of MGv, all of which have frequency-related topography (Horie et al., 2013; Tsukano et al., 2015, 2017a, 2017b). IAF in the insular cortex also receives topological projections from the ventro-medial part of middle MGv (Takemoto et al., 2014). Finally, the current work revealed that A2 receives topological projections from neurons in the caudal part of MGv. These facts suggest that the five topological thalamocortical pathways convey some specific sound information whose cortical target is selected by gating in MGv (Blundon and Zakharenko, 2013).

Identifying the thalamic origins of A2 makes homologizing regions between animals easier. In mice, A2 sits the most ventrally in the auditory cortex (Figure 1), its neurons have the longest response latency (Guo et al., 2012; Kubota et al., 2008), and MGv neurons projecting to A2 are localized caudally (Figure 3). A similar fine-grained map and thalamocortical connections have been shown in rats (Storace et al., 2010; Polley et al., 2007). In rats, the suprarhinal auditory field (SRAF) is equivalent to mouse A2, being the most ventral tonotopic region with long response latency, and thalamic neurons projecting to SRAF are localized caudally in MGv (Storace et al., 2012). These common features between mouse A2 and rat SRAF indicate that they are homologous. Considering the presence of parallel thalamocortical inputs in cats, a common processing principle for mammalian audition might involve neuronal computations that integrate distinct thalamocortical and corticocortical inputs.

### Thalamocortical projection-dependent mechanisms for differentiating frequency organization in the auditory system

Two-photon imaging studies indicate that increased heterogeneity in BF distribution and broader tuning curves follow along the hierarchical streams in the auditory cortex (Issa et al., 2014; Winkowski and Kanold, 2013). Increased tonotopic heterogeneity can be explained by the large divergence we found in projections from MGv to A2, which could lead to the recently found heterogeneous BF distribution in presynaptic thalamocortical buttons (Vasquez-Lopez et al., 2017). The consistency between the large MGv-A2 divergence and the wide tuning curves in A2 can be explained by the fact that tuning curves in the auditory cortex are generally determined by thalamic inputs over intracortical circuits (Liu et al., 2007). Additionally, the relatively ordered tonotopy in the thalamocortical input layer of A1 can be explained by the small divergence in MGv-A1 projections. Overall, the current results demonstrate the presence of a complexity generator in inputs to A2, which exemplifies the concept of thalamocortical transformation of functional organization due to divergence in thalamocortical projections (Imaizumi and Lee, 2014). Moreover, A2 receives about twice as many thalamic projections from MGd as does A1 (Figure 3A) (Takemoto et al., 2014), and they synapse in the same layer that receives input from MGv (Llano and Sherman, 2008). MGd neurons exhibit complex response patterns such as bursting in response to tones (He and Hu, 2002) and broad bandwidth (Calford, 1983), therefore summation of divergent inputs from MGv and inputs with complex responses from MGd likely leads to more complex output from neurons in thalamocortical recipient layer of A2. Again, transformation of the local functional organization according to the hierarchy is not spontaneous but can be realized by an anatomical mechanism in the thalamocortical ascending pathways. Such a structure-based, complexity generator might be the foundation for realizing higher-order functions in A2 beyond tonotopy such as recognizing biological values of a given sound (Geissler et al., 2016).

## Materials and methods

### Animals

This study was conducted in accordance with the approved protocols and guidelines of the Committee for Animal Care in Niigata University. We used 6–7-week old male C57BL/6N mice (Charles River Japan, Kanagawa, Japan). Mice were housed in cages under a 12 hours light/dark cycle with *ad libitum* access to food pellets and water.

### Flavoprotein fluorescence imaging

Flavoprotein fluorescence imaging was performed according to methods in our previous studies (Horie et al., 2013; Tsukano et al., 2015). Mice were anesthetized with urethane (1.65 g/kg, i.p.) and rectal temperature was kept at ~37.0 °C. After subcutaneous injection of bupivacaine, the skin and temporal muscle were removed. A piece of metal was attached to the skull with dental resin, and the head was fixed by screwing the metal piece onto a manipulator. Unless otherwise noticed, a ~3 × 3-mm craniotomy was performed over the right auditory cortex. For transcranial imaging (Figure 1), the skull over the right auditory cortex was intact and kept transparent using liquid paraffin. Cortical images (128 × 168 pixels after binning) of endogenous green fluorescence (λ = 500–550 nm) in blue light (λ = 470–490 nm) were recorded at 9 Hz using a cooled charge-coupled device (CCD) camera system (AQUACOSMOS with ORCA-R2 camera; Hamamatsu Photonics, Shizuoka, Japan) that was attached to an epifluorescence microscope (M651; Leica, Wetzlar, Germany). Images were averaged over 20 trials. Fluorescence responses were normalized as ΔF/F_0_, where F_0_ represents the average of five images (~500 ms) before stimulus onset.

### Neural tracer injections

We used Fast Blue (Funakoshi, Tokyo, Japan), Alexa Fluor-488, and Alexa Fluor-555 conjugated CTB (Invitrogen, Thermo Fisher Scientific, Boston, MA) as a retrograde neural tracer. We performed triple injections of Fast Blue, CTB-488, and CTB-555, or dual injections of CTB-488 and CTB-555. Before injection, precise locations of auditory regions were identified using flavoprotein fluorescence imaging. A glass pipette (tip: 20–30 μm) filled with a tracer was inserted ~400 μm down into an identified area. One of the tracers was then injected iontophoretically with a 5-μA anodal currents (5 s on, 5 s off) having ~100 pulses. After the injection, the glass pipette was slowly withdrawn and the skin was sutured. Mice were placed in a warm place for recovery, and after awaking they were reared in their home cages.

### Tracer observation

Three days after injection, mice were anesthetized with an overdose of pentobarbital (0.3 g/kg), and cardiac perfusion was performed using 4% paraformaldehyde. After sequential immersion into a 20% and 30% sucrose solution, coronal or horizontal brain slices were prepared at 40-μm intervals according to our previous studies (Horie et al., 2013, 2015). Every fourth slice was mounted on glass slides and cover-slipped using Fluoromount (Cosmo Bio, Tokyo, Japan). Sections were observed using a light microscope (Eclipse Ni, Nikon, Tokyo, Japan) and a CCD camera (DP-80; Olympus, Tokyo, Japan). After observing fluorescent tracers, sections were Nissl-stained as described in Horie et al. 2013, and cover-slipped using Bioleit (Okenshoji, Tokyo, Japan).

### Suction injury

After identifying the locations of auditory cortices using optical imaging, we lesioned the auditory cortex by slowly sucking all layers using a glass pipette (tip diameter: ~70 μm) connected to a vacuum pump. Suction was stopped at the depth where the parenchyma turned whitish in color, which indicated the white matter beneath the auditory cortex had been reached. After completing the suction, the hole was covered with 2% agarose (1-B, Sigma-Aldrich, MO) and a thin cover glass (thickness <0.15 mm, Matsunami, Osaka, Japan), which was fixed to the skull with dental cement (Sunmedical, Siga, Japan).

### Partition of the MGB subdivisions

The borders of the medial geniculate body (MGB) subdivisions were drawn according to the mouse brain atlas (Paxinos and Franklin, 2012) and SMI-32 immunolabeling patterns, which recognize a non-phosphorylated epitope on the 168- and 200-kDa subunits of neurofilament proteins (Horie et al., 2013, 2015; LeDoux et al., 1985, 1987; Honma et al., 2013). SMI-32 immunolabeling was performed in adjacent sections. Sections were rinsed and incubated in PBST containing 3% hydrogen peroxide. After rinsing in 20 mM PBS, the sections were incubated overnight at room temperature with the monoclonal antibody SMI-32 (1:2000, Covance Research Products, Berkeley, CA), diluted with 20 mM PBS containing 0.5% skim milk. Sections were then incubated with horseradish peroxidase (HRP)-labeled anti-mouse IgG (1:100, MBL, Nagoya, Japan) at room temperature for 2 h. The sections were then rinsed in 20 mM PBS, and the immunoreactions was visualized in a Tris-HCl buffer containing 0.05% diaminobenzidine tetrahydrochloride and 0.003% hydrogen peroxide for 4–5 minutes. In some slices, we conducted calbindin immunolabeling because MGd exhibits strong immunoreactivity to calbindin (Takemoto et al., 2014; Lu et al., 2009). We used anti-calbindin primary antibody (Abcam, Cambridge, UK) and HRP-labeled anti-Rabbit IgG as a secondary antibody (Invitrogen Life Technologies, Boston, MA). The remaining protocols were the same as those used to visualize SMI-32.

### Separation Index

To quantify and compare the degree of spatial overlap between two neuronal populations, we used the absolute value of Cliff’s delta (Cliff, 1993) as a Separation Index (SI). Cliff’s delta is a non-parametric metric of an effect size described in the following formula,

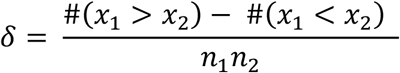

where *n_i_* is the size of group *i* (*i* = 1, 2) and #(*x_i_* > *x_j_*) is an operator that counts the number of all possible combinations such that an item of group 1 is greater than that of group 2. It follows that Cliff’s delta ranges from −1 to 1 and the three extreme conditions are

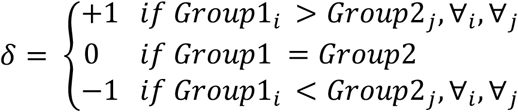

Because we did not need to discriminate the direction of difference in our analysis, we defined the absolute value of Cliff’s delta as SI. The value ranges from 0 to 1, with 0 indicating that the locations of the two groups completely overlap and 1 indicating that they are completely separate without any overlap.

### *In vivo* two-photon calcium imaging

After inducing anesthesia with urethane (1.9 g/kg, i.p.) and craniotomy of the right auditory cortex, the locations of A1 and A2 were identified using flavoprotein fluorescence imaging. Calcium imaging was performed using a two-photon microscope (TCS SP5 MP, Leica Microsystems, Wetzlar, Germany) with a hybrid detector (HyD, Leica Microsystems) and a Ti-Sapphire mode-locked femtosecond laser (Chameleon Vision, Coherent, Santa Clara, CA). Images were obtained via a ×20 water-immersion objective lens (numerical aperture 1.0, HCX PL APO; Leica Microsystems). We used Cal-520 as a calcium indicator because it exhibits high S/N ratio and its event-related fluorescence change is strongly correlated with spike number (Li et al., 2017; Tada et al., 2014). Cal-520 AM (Invitrogen, Thermo Fisher Scientific) was dissolved in 10% (w/v) Pluronic F-127 in DMSO and diluted with Ringer solution containing sulforhodamine 101 (SR-101, Invitrogen, Thermo Fisher Scientific). The Cal-520 solution was pressure injected (5–20 kPa) for 5–10 min into layer 3b/4 using glass pipettes (tip diameter: 2–4 μm). Astrocytes were distinguished from neurons using SR-101. After injection, the pipette was withdrawn and the craniotomy was covered with 2% agarose (1-B, Sigma-Aldrich) and a thin cover glass (thickness <0.15 mm, Matsunami), which was fixed to the skull with dental cement (Sunmedical). Excitation wavelength for Cal-520 was 900 nm and that for SR-101 was 940–950 nm. Cal-520 fluorescence was observed between 500 and 550 nm. Images (256 × 256 pixels) were recorded at 3.7 Hz in a 246 × 246-μm region. Rectal temperature of mice was kept at ~37.0 °C.

Data were realigned using AQUACOSMOS and custom-written MATLAB code (Mathworks, St. Louise, MO). Data from 5–6 repeated trials of the same stimulation were averaged. Size-matched ROIs were chosen. Data were calculated as ΔF/F_0_, where ΔF = F – F_0_, and F_0_ (the baseline intensity) was obtained by averaging the intensity values during the pre-stimulus period (about 3 s) across the repeated trials. The response of each neuron was evaluated in poststimulus observation windows (about 5 s). The BF of a neuron was defined as the frequency to which the neuron had the greatest response. We examined the correlation between BF and location and computed a regression line. The direction of frequency organization was defined so as to make the correlation coefficient the largest. Bandwidth of the tuning curves was defined as the logarithmic ratio of minimum and maximum frequencies that resulted in a response >75% of the peak amplitude.

### Auditory stimuli

Tones were generated by a computer using a custom-written LabVIEW code (National Instruments, Austin, TX). The sampling rate was 500 kHz. Sounds were low-pass filtered at 150 kHz (3624, NF, Kanagawa, Japan). Pure tones at frequencies of 5–80 kHz were amplitude modulated by a 20-Hz sine wave. A speaker for 5–40-kHz (SRS-3050A, Stax, Saitama, Japan) or 50–80-kHz (ES105A, Murata, Kyoto, Japan) tones was set 10 cm in front of the mice. Sound intensity was calibrated using a microphone (Type 4135 and Type 2669, Brüel & Kjær, Nærum, Denmark) and a sound-level meter (Type 2610, Brüel & Kjær). The sound intensity was set to 60 dB SPL, and the duration was 500 ms with a rise/fall time of 10 ms. The desired sound spectrum was confirmed using a digital spectrum analyzer (R9211A, Advantest, Tokyo, Japan) or a custom-written LabVIEW program.

### Statistics

Statistical analyses were conducted using MATLAB programs, and SPSS (IBM Corp, Armonk, NY) if necessary. The Mann–Whitney U-test or the Wilcoxon signed-rank test was used to evaluate differences between two paired or unpaired two groups, respectively. The *χ*^2^ test was used to evaluate differences in proportions of neurons. Correlation was evaluated using Spearman’s test. The Kormogorv–Smirnov test was used to evaluate the differences between two cumulative distributions. When multiple comparisons were needed, we used Bonferroni correction. We evaluated the differences between two correlation coefficients according to Howell, 2012. Briefly, two r values were *z*-transformed by the function *z* = *atanh*(*r*) and the *Z* test statistic was calculated as

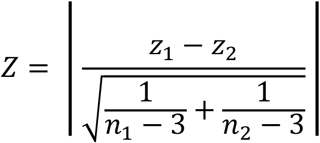

P-values were obtained by the formula,

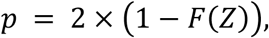

where *F*(*Z*) is the normal cumulative distribution function of *Z*.

If MATLAB provided p-values of zero in any statistical analysis, we replaced them with p < 10^−35^, because null hypotheses cannot be rejected with a probability of 100%.

## Acknowledgements

This work was supported by JSPS KAKENHI Grant No. JP17K07051 (to H. Tsukano) and JP26830008 (to H. Tsukano), a grant for the Promotion of Medical Science and Medical Care No. 15KI149 from the Ichiro Kanehara Foundation (to H. Tsukano), and a grant for Basic Science Research Projects No. 140254 from the Sumitomo Foundation (to H. Tsukano). We thank Saeko Maruyama for technical assistance, and Ayano Matsushima and Mari Isogai for animal breeding and maintenance. We also thank Dr. Adam Phillips from Edanz Group (www.edanzediting.com/ac) for editing a draft of this manuscript.

## Author Contributions

S.O., H. Tsukano, and M.H. performed experiments. S.O., H. Tsukano, M.H., H. Terashima, and N.N. analyzed the data. Y.K. and K.T. provided critical ideas for planning this work. R.H. and H. Takebayashi provided critical comments regarding the results. S.O., H. Tsukano, and M.H. prepared the Figures. H. Tsukano wrote the manuscript. H. Tsukano and K.S. revised the manuscript. All authors approved the publication of the manuscript.

## Declaration of Interests

The authors declare no conflict of interest.

## References

Bandyopadhyay S, Shamma SA, Kanold PO (2010) Dichotomy of functional organization in the mouse auditory cortex. Nat Neurosci 13:361–368.

Bizley JK, King AJ (2009) Visual influences on ferret auditory cortex. Hear Res 258:55–63.

Blundon JA, Zakharenko SS (2013) Presynaptic gating of postsynaptic synaptic plasticity: a plasticity filter in the adult auditory cortex. Neuroscientist 19:465–478.

Calford MB (1983) The parcellation of the medial geniculate body of the cat defined by the auditory response properties of single units. J Neurosci 3:2350–2364.

Castro JB, Kandler K (2010) Changing tune in auditory cortex. Nat Neurosci 13:271–273.

Chen JL, Carta S, Soldado-Magraner J, Schneider BL, Helmchen F (2013) Behaviour-dependent recruitment of long-range projection neurons in somatosensory cortex. Nature 499:336–340.

Cliff N (1993) Dominance statistics: Ordinal analyses to answer ordinal questions. Psychol Bull 114:494–509.

Covic EN, Sherman SM (2011) Synaptic properties of connections between the primary and secondary auditory cortices in mice. Cereb Cortex 21:2425–2441.

de la Mothe LA, Blumell S, Kajikawa Y, Hackett TA (2012) Thalamic connections of auditory cortex in marmoset monkeys: lateral belt and parabelt regions. Anat Rec 295:822–836.

Gazzaniga MS (2000) Cerebral specialization and interhemispheric communication: does the corpus callosum enable the human condition? Brain 123:1293–1326.

Geissler DB, Schmidt HS, Ehret G (2016) Knowledge About Sounds-Context-Specific Meaning Differently Activates Cortical Hemispheres, Auditory Cortical Fields, and Layers in House Mice. Front Neurosci 10:98.

Glickfeld LL, Andermann ML, Bonin V, Reid RC (2013) Cortico-cortical projections in mouse visual cortex are functionally target specific. Nat Neurosci 16:219–226.

Guo W, Chambers AR, Darrow KN, Hancock KE, Shinn-Cunningham BG, Polley DB (2012) Robustness of cortical topography across fields, laminae, anesthetic states, and neurophysiological signal types. J Neurosci 32:9159–9172.

Harel N, Mori N, Sawada S, Mount RJ, Harrison RV (2000) Three distinct auditory areas of cortex (A1, A2, and AAF) defined by optical imaging of intrinsic signals. Neuroimage 11:302–312.

He J, Hu B (2002) Differential distribution of burst and single-spike responses in auditory thalamus. J Neurophysiol 88:2152–2256.

Higgins NC, Storace DA, Escabí MA, Read HL (2010) Specialization of binaural responses in ventral auditory cortices. J Neurosci 30:14522–14532.

Honma Y, Tsukano H, Horie M, Ohshima S, Tohmi M, Kubota Y, Takahashi K, Hishida R, Takahashi S, Shibuki K (2013) Auditory cortical areas activated by slow frequency-modulated sounds in mice. PLoS One 8:e68113.

Horie M, Tsukano H, Hishida R, Takebayashi H, Shibuki K (2013) Dual compartments of the ventral division of the medial geniculate body projecting to the core region of the auditory cortex in C57BL/6 mice. Neurosci Res 76:207–212.

Horie M, Tsukano H, Takebayashi H, Shibuki K (2015) Specific distribution of non-phosphorylated neurofilaments characterizing each subfield in the mouse auditory cortex. Neurosci Lett 606:182–187.

Howell DC (2012) Correlation and Regression. Statistical Methods for Psychology, 8th ed, eds Howell DC (Wadsworth, Belmont) pp 251–301.

Huang CL, Winer JA (2000) Auditory thalamocortical projections in the cat: laminar and areal patterns of input. J Comp Neurol 427:302–331.

Imaizumi K, Lee CC (2014) Frequency transformation in the auditory lemniscal thalamocortical system. Front Neural Circuits 8:75.

Issa JB, Haeffele BD, Agarwal A, Bergles DE, Young ED, Yue DT (2014) Multiscale Optical Ca2+ Imaging of Tonal Organization in Mouse Auditory Cortex. Neuron 83:944–959.

Joachimsthaler B, Uhlmann M, Miller F, Ehret G, Kurt S (2014) Quantitative analysis of neuronal response properties in primary and higher-order auditory cortical fields of awake house mice (Mus musculus). Eur J Neurosci 39:904–918.

Kaas JH, Hackett TA (2000) Subdivisions of auditory cortex and processing streams in primates. Proc Natl Acad Sci USA 97:11793–11799.

Kalatsky VA, Polley DB, Merzenich MM, Schreiner CE, Stryker MP (2005) Fine functional organization of auditory cortex revealed by Fourier optical imaging. Proc Natl Acad Sci USA 102:13325–13330.

Kato HK, Gillet SN, Isaacson JS (2015) Flexible Sensory Representations in Auditory Cortex Driven by Behavioral Relevance. Neuron 88:1027–1039.

Kubota Y, Kamatani D, Tsukano H, Ohshima S, Takahashi K, Hishida R, Kudoh M, Takahashi S, Shibuki K (2008) Transcranial photo-inactivation of neural activities in the mouse auditory cortex. Neurosci Res 60:422–430.

LeDoux JE, Ruggiero DA, Forest R, Stornetta R, Reis DJ (1987) Topographic organization of convergent projections to the thalamus from the inferior colliculus and spinal cord in the rat. J Comp Neurol 264:123–146.

LeDoux JE, Ruggiero DA, Reis DJ (1985) Projections to the subcortical forebrain from anatomically defined regions of the medial geniculate body in the rat. J Comp Neurol 242:182–213.

Lee CC (2015) Exploring functions for the non-lemniscal auditory thalamus. Front Neural Circuits 9:69.

Lee CC, Schreiner CE, Imaizumi K, Winer JA (2004) Tonotopic and heterotopic projection systems in physiologically defined auditory cortex. Neuroscience 128:871–887.

Lee CC, Sherman SM (2008) Synaptic properties of thalamic and intracortical inputs to layer 4 of the first- and higher-order cortical areas in the auditory and somatosensory systems. J Neurophysiol 100:317–326.

Lee CC, Sherman SM (2010) Drivers and modulators in the central auditory pathways. Front Neurosci 4:79–86.

Lee CC, Sherman SM (2010) Topography and physiology of ascending streams in the auditory tectothalamic pathway. Proc Natl Acad Sci USA 107:372–377.

Lee CC, Winer JA (2008) Connections of cat auditory cortex: I. Thalamocortical system. J Comp Neurol 507:1879–1900.

Li J, Zhang J, Wang M, Pan J, Chen X, Liao X (2017) Functional imaging of neuronal activity of auditory cortex by using Cal-520 in anesthetized and awake mice. Biomed Opt Express 8:2599–2610.

Liu BH, Wu GK, Arbuckle R, Tao HW, Zhang LI (2007) Defining cortical frequency tuning with recurrent excitatory circuitry. Nat Neurosci 10:1594–1600.

Llano DA, Sherman SM (2008) Evidence for nonreciprocal organization of the mouse auditory thalamocortical-corticothalamic projection systems. J Comp Neurol 507:1209–1227.

Lu E, Llano DA, Sherman SM (2009) Different distributions of calbindin and calretinin immunostaining across the medial and dorsal divisions of the mouse medial geniculate body. Hear Res 257:16–23.

Nishimura M, Shirasawa H, Kaizo H, Song WJ (2007) New field with tonotopic organization in guinea pig auditory cortex. J Neurophysiol 97:927–932.

Oviedo HV, Bureau I, Svoboda K, Zador AM (2010) The functional asymmetry of auditory cortex is reflected in the organization of local cortical circuits. Nat Neurosci 13:1413–1420.

Paxinos G, Franklin, KBJ (2012) The Mouse BrA1n in Stereotaxic Coordinates, Fourth Edition. San Diego, CA: Academic Press.

Petkov CI, Kayser C, Augath M, Logothetis NK (2006) Functional imaging reveals numerous fields in the monkey auditory cortex. PLoS Biol 4:e215.

Polley DB, Read HL, Storace DA, Merzenich MM (2007) Multiparametric auditory receptive field organization across five cortical fields in the albino rat. J Neurophysiol 97:3621–3638.

Rothschild G, Nelken I, Mizrahi A (2010) Functional organization and population dynamics in the mouse primary auditory cortex. Nat Neurosci 13:353–360.

Saenz M, Langers DR (2014) Tonotopic mapping of human auditory cortex. Hear Res 307:42–52.

Shiramatsu TI, Takahashi K, Noda T, Kanzaki R, Nakahara H, Takahashi H (2016) Microelectrode mapping of tonotopic, laminar and field-specific organization of thalamo-cortical pathway in rat. Neuroscience 332:38–52.

Smith PH, Uhlrich DJ, Manning KA, Banks MI (2012) Thalamocortical projections to rat auditory cortex from the ventral and dorsal divisions of the medial geniculate nucleus. J Comp Neurol 520:34–51.

Stiebler I, Neulist R, Fichtel I, Ehret G (1997) The auditory cortex of the house mouse: left-right differences, tonotopic organization and quantitative analysis of frequency representation. J Comp Physiol A 181: 559–571.

Storace DA, Higgins NC, Chikar JA, Oliver DL, Read HL (2012) Gene expression identifies distinct ascending glutamatergic pathways to frequency-organized auditory cortex in the rat brain. J Neurosci 32:15759–15768.

Storace DA, Higgins NC, Read HL (2010) Thalamic label patterns suggest primary and ventral auditory fields are distinct core regions. J Comp Neurol 518:1630–1646.

Storace DA, Higgins NC, Read HL (2011) Thalamocortical pathway specialization for sound frequency resolution. J Comp Neurol 519:177–193.

Tada M, Takeuchi A, Hashizume M, Kitamura K, Kano M (2014) A highly sensitive fluorescent indicator dye for calcium imaging of neural activity in vitro and in vivo. Eur J Neurosci 39:1720–1728.

Takemoto M, Hasegawa K, Nishimura M, Song WJ (2014) The insular auditory field receives input from the lemniscal subdivision of the auditory thalamus in mice. J Comp Neurol 522:1373–1389.

Tao C, Zhang G, Zhou C, Wang L, Yan S, Tao HW, Zhang LI, Zhou Y, Xiong Y (2017) Diversity in Excitation-Inhibition Mismatch Underlies Local Functional Heterogeneity in the Rat Auditory Cortex. Cell Rep 19:521–531.

Tsukano H, Horie M, Bo T, Uchimura A, Hishida R, Kudoh M, Takahashi K, Takebayashi H, Shibuki K (2015) Delineation of a frequency-organized region isolated from the mouse primary auditory cortex. J Neurophysiol 113:2900–2920.

Tsukano H, Horie M, Hishida R, Takahashi K, Takebayashi H, Shibuki K (2016) Quantitative map of multiple auditory cortical regions with a stereotaxic fine-scale atlas of the mouse brain. Sci Rep 6:22315.

Tsukano H, Horie M, Ohga S, Takahashi K, Kubota Y, Hishida R, Takebayashi H Shibuki K (2017) Reconsidering Tonotopic Maps in the Auditory Cortex and Lemniscal Auditory Thalamus in Mice. Front Neural Circuits 11:14.

Tsukano H, Horie M, Takahashi K, Hishida R, Takebayashi H Shibuki K (2017) Independent tonotopy and thalamocortical projection patterns in two adjacent parts of the classical primary auditory cortex in mice. Neurosci Lett 637:26–30.

Vasquez-Lopez SA, Weissenberger Y, Lohse M, Keating P, King AJ, Dahmen JC (2017) Thalamic input to auditory cortex is locally heterogeneous but globally tonotopic. eLife in press.

Wang Q, Burkhalter A (2007) Area map of mouse visual cortex. J Comp Neurol 502:339–357.

Winer JA, Larue DT, Huang CL (1999) Two systems of giant axon terminals in the cat medial geniculate body: convergence of cortical and GABAergic inputs. J Comp Neurol 413:181–197.

Winkowski DE, Kanold PO (2013) Laminar transformation of frequency organization in auditory cortex. J Neurosci 33:1498–1508.

Zhang G, Tao C, Zhou C, Yan S, Wang Z, Zhou Y, Xiong Y (2017) Excitatory effects of the primary auditory cortex on the sound-evoked responses in the ipsilateral anterior auditory field in rat. Neuroscience 361:157–166.

Zhou M, Liang F, Xiong XR, Li L, Li H, Xiao Z, Tao HW, Zhang LI (2014) Scaling down of balanced excitation and inhibition by active behavioral states in auditory cortex. Nat Neurosci 17:841–850.

